# NextSV: a meta-caller for structural variants from low-coverage long-read sequencing data

**DOI:** 10.1101/092544

**Authors:** Li Fang, Jiang Hu, Depeng Wang, Kai Wang

## Abstract

**Background:** Structural variants (SVs) in human genomes are implicated in a variety of human diseases. Long-read sequencing delivers much longer read lengths than short-read sequencing and may greatly improve SV detection. However, due to the relatively high cost of long-read sequencing, it is unclear what coverage is needed and how to optimally use the aligners and SV callers.

**Results:** In this study, we developed NextSV, a meta-caller to perform SV calling from low coverage long-read sequencing data. NextSV integrates three aligners and three SV callers and generates two integrated call sets (sensitive/stringent) for different analysis purposes. We evaluated SV calling performance of NextSV under different PacBio coverages on two personal genomes, NA12878 and HX1. Our results showed that, compared with running any single SV caller, NextSV stringent call set had higher precision and balanced accuracy (F1 score) while NextSV sensitive call set had a higher recall. At 10X coverage, the recall of NextSV sensitive call set was 93.5% to 94.1% for deletions and 87.9% to 93.2% for insertions, indicating that ~10X coverage might be an optimal coverage to use in practice, considering the balance between the sequencing costs and the recall rates. We further evaluated the Mendelian errors on an Ashkenazi Jewish trio dataset.

**Conclusions:** Our results provide useful guidelines for SV detection from low coverage whole-genome PacBio data and we expect that NextSV will facilitate the analysis of SVs on long-read sequencing data.

## Background

Structural variants (SVs) represent genomic rearrangements (typically defined as longer than 50 bp), and SVs may play important roles in human diversity and disease susceptibility [1–3]. Many inherited diseases and cancers have been associated with a large number of SVs in recent years [4–9]. Recent advances in next-generation sequencing (NGS) technologies have facilitated the analysis of variations such as SNPs and small indels in unprecedented details, but the discovery of SVs using short-read sequencing still remains challenging [10]. Single-molecule, real-time (SMRT) sequencing developed by Pacific Biosciences (PacBio) produces long-read sequencing data, making it potentially well-suited for SV detection in personal genomes [10, 11]. Most recently, Merker et al. reported the application of low coverage whole genome PacBio sequencing to identify pathogenic structural variants from a patient with autosomal dominant Carney complex, for whom targeted clinical gene testing and whole genome short-read sequencing were both negative [12]. This represents a clear example that long-read sequencing may solve some negative cases in clinical diagnostic settings.

Two popular SV software tools have been developed specifically for long-read sequencing: PBHoney [13] and Sniffles [14]. PBHoney identifies genomic variants via two algorithms, long-read discordance (PBHoney-Spots) and interrupted mapping (PBHoney-Tails). Sniffles is a SV caller written in C++ and it detects SVs using evidence from split-read alignments, high-mismatch regions, and coverage analysis [14]. PBHoney uses BAM files generated by BLASR [15] as input while Sniffles requires BAM files from BWA-MEM [16] or NGMLR [14], a new long-read aligner. Due to the relatively high cost of PacBio sequencing, users are often faced with issues such as what coverage is needed and how to get the best use of the available aligners and SV callers. In addition, it is unclear which software tool performs the best in low-coverage settings, and whether the combination of software tools can improve performance of SV calls. Finally, the execution of these software tools is often not straightforward and requires careful re-parameterization given specific coverage of the source data.

To address these challenges, we developed NextSV, an automated SV detection pipeline integrating multiple tools. NextSV automatically execute these software tools with optimized parameters for user-specified coverage, then integrates results of each caller and generates a sensitive call set and a stringent call set, for different analysis purposes.

Recently, the Genome in a Bottle (GIAB) consortium and the 1000 Genome Project Consortium released high-confidence SV calls for the NA12878 genome, an extensively sequenced genome by different platforms, enabling benchmarking of SV callers [17, 18]. They also published sequencing data of seven human genomes, including PacBio data of an Ashkenazi Jewish (AJ) family trio [19]. Previously, we sequenced a Chinese individual HX1 on the PacBio platform with over 100X coverage, and generated assembly-based SV call sets [20]. Using data sets of NA12878, HX1 and the AJ family trio, we evaluated the performance of four aligner/SV caller combinations (BLASR / PBHoney-Spots, BLASR / PBHoney-Tails, BWA / Sniffles and NGMLR / Sniffles) as well as NextSV under different PacBio coverages. We expect that NextSV will facilitate the detection and analysis of SVs on long-read sequencing data.

## Materials and Methods

### PacBio data sets used for this study

Five whole-genome PacBio sequencing data sets were used to test the performance of SV calling pipelines (Table 1). Data sets of NA12878 and HX1 genome were downloaded from NCBI SRA database (Accession: SRX627421, SRX1424851). Data sets of the AJ family trio were downloaded from the FTP site of National Institute of Standards and Technology (NIST) [21]. After we obtained raw data, we extracted subreads (reads that can be used for analysis) using the SMRT Portal software (Pacific Biosciences, Menlo Park, CA) with filtering parameters (minReadScore=0.75, minLength=500). The subreads were mapped to the reference genome using BLASR [15], BWA-MEM [16] or NGMLR [14]. The BAM files were down-sampled to different coverages using SAMtools (samtools view -s). We performed five subsampling replicates at each coverage. The down-sampled coverages and mean read lengths of the data sets were shown in Table 1.

**Table 1.** Description of PacBio data sets used for this study.

| Data Source | Genome | Original Coverage | Down-sampled Coverage | Mean Read Length | Reference |
| --- | --- | --- | --- | --- | --- |
| NCBI SRA | NA12878 | 22X | 2-15X | 4.9 kb | [26] |
| NCBI SRA | HX1 | 103X | 6-15X | 7.0 kb | [20] |
| NIST | AJ son | 69X | 10X | 8.0 kb | [19] |
| NIST | AJ father | 32X | 10X | 7.3 kb | [19] |
| NIST | AJ mother | 30X | 10X | 7.8 kb | [19] |

### SV detection using BLASR / PBHoney-Spots and BLASR / PBHoney-Tails

PacBio subreads were iteratively aligned to the human reference genome (GRCh38 for HX1, GRCh37 for NA12878 and AJ trio genomes, depending on the reference of high-confidence set) using the BLASR aligner (parameter: -bestn 1). Each read’s single best alignment was stored in the SAM output. Unmapped portions of each read were extracted from the alignments and remapped to the reference genome. The alignments in SAM format were converted to BAM format and sorted by SAMtools. PBHoney-Tails and PBHoney-Spots (from PBSuite-15.8.24) were run with slightly modified parameters (minimal read support 2, instead of 3 and consensus polishing disabled) to increase sensitivity and to discover SVs under low coverages (2–15X). The reference FASTA files used in this study were downloaded from the FTP sites of 1000 Genome Project [22] (GRCh37) and NCBI [23] (GRCh38). The FASTA files contain assembled chromosomes with unlocalized, unplaced and decoy sequences.

### SV detection using BWA / Sniffles and NGMLR / Sniffles

PacBio subreads were aligned to the reference genome, using BWA-MEM (bwa mem -M -x pacbio) or NGMLR (default parameters) to generate the BAM file. The BAM file was sorted by SAMtools, then used as input of Sniffles (version 1.0.5). Sniffles was run with slightly modified parameters (minimal read support 2, instead of 10) to increase sensitivity and discover SVs under low fold of coverages (2–15X).

### NextSV analysis pipeline

As shown in Figure 1, NextSV currently supports four aligner / SV caller combinations: BLASR / PBHoney-Spots, BLASR / PBHoney-Tails, BWA / Sniffles and NGMLR / Sniffles. NextSV extracts FASTQ files from PacBio raw data (.hdf5 or .bam) and performs QC according to users specified settings. Once the aligner / SV caller combination is selected by user, NextSV automatically generates the scripts for alignment, sorting, and SV calling with appropriate parameters. When the analysis is finished, NextSV will format the raw result files (.tails, .spots, or .vcf files) into BED files. If multiple aligner/SV caller combinations are selected, NextSV will integrate the calls to generate a sensitive (by union) and a stringent (by intersection) call set. The output of NextSV is ANNOVAR-compatible, so that users can easily perform downstream annotation using ANNOVAR [24]. In addition, NextSV also supports job submitting via Sun Grid Engine (SGE), a popular batch-queuing system in cluster environment.

**Figure 1.**
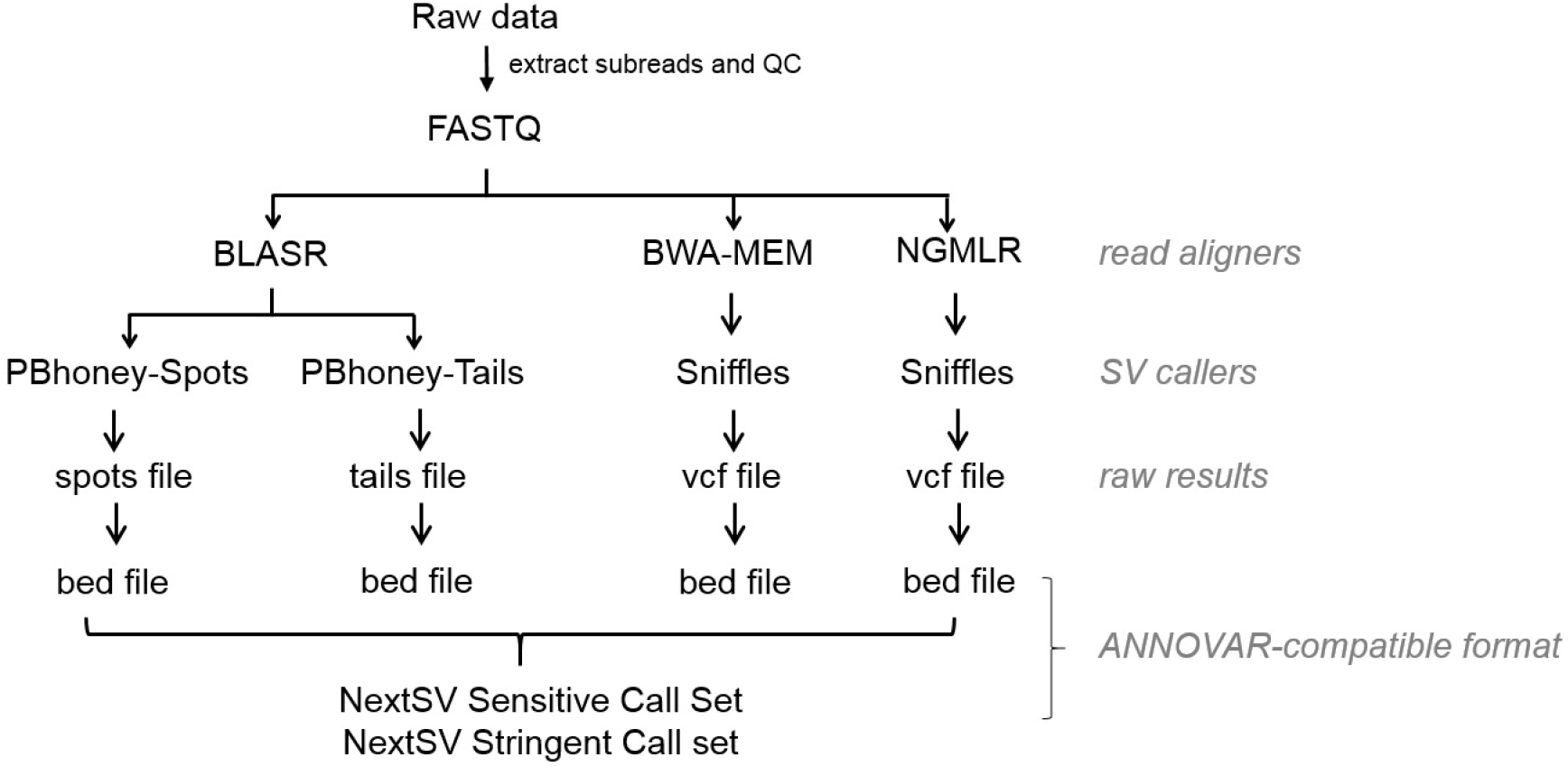
Scheme of NextSV workflow.

Users can choose to run any of the four aligner/SV caller combination. By default, NextSV will enable BLASR / PBHoney-Spots, BLASR / PBHoney-Tails and NGMLR / Sniffles and integrate the results to generate the sensitive calls and stringent calls. We do not enable BWA / Sniffles by default because Sniffles works better with NGMLR in our evaluation and alignment is a time consuming step. SVs that are shorter than reads may result in intra-read discordances while larger SVs may result in soft-clipped tails of long reads. We suggest running both PBHoney-Spots and PBHoney-Tails because they are two complementary algorithms designed to detect intra-read discordances and soft-clipped tails, respectively. Sniffles uses multiple evidences to detect SV so it should be suitable for both types of SVs.

NextSV sensitive call set is generated as:

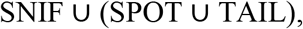
and NextSV stringent call set is generated as:

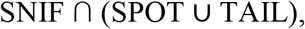
where SNIF denotes the call set of Sniffles (the aligner can be BWA or NGMLR, whichever is enabled; if both aligners are enabled, the call set of NGMLR/Sniffles will be used), SPOT denotes the call set of BLASR / PBHoney-Spots and TAIL denotes the call set of BLASR / PBHoney-Tails.

### Comparing two SV call sets

The criteria for merging two SV calls were chosen to follow what was done by the NIST/GIAB analysis team [25] and a previous study [26]. Two deletion calls were considered the same if they had at least 50% reciprocal overlap (the overlapped region was more than 50% of both calls). The insertion call had a single breakpoint position so the criterion for insertion calls should be different from that of deletion calls. Two insertion calls were considered the same if the two breakpoints were within a distance *delta. Delta* used by NIST/GIAB analysis team was 1000 bp and used by Pendleton et al (reference [26]) was 100 bp. However, 100 bp was too small for our analysis since the coverages (2–15X) were far lower than that of Pendleton’s data set (46X in total). On the other hand, 1000 bp might be too large to include distant calls as the same merged call. Therefore, we chose 500 bp as a compromise. When merging two SVs, the average start and end positions were taken.

### High-confidence SV call sets

The high-confidence deletion call set of the NA12878 genome was release by the Genome In A Bottle (GIAB) consortium [17], in which most of the calls were refined by experimental validation or other independent technologies. The high-confidence insertion call set of the NA12878 genome was obtained by merging the high-confidence insertion calls of 1000 Genome phase 3 [18] and high-confidence insertion calls from GIAB. For the HX1 genome, we generated the high-confidence SV call set via two steps. First, we used the SV calls from a previously validated local assembly-based approach [11] as the initial high-quality calls. Next, we detected SVs on 103X coverage PacBio data set of the HX1 genome using BLASR / PBHoney-Spots, BLASR / PBHoney-Tails, BWA / Sniffles and NGMLR / Sniffles (minimal read support=20 for each SV caller). The initial high-quality calls (from step 1) that overlapped with one of the four 103X call sets (from step 2) were retained as final high-confidence calls. SVs are generally defined as genomic rearrangements that are larger than 50 bp. However, we do not consider SVs that are less than 200 bp. There are two reasons. First, SVs that are smaller than 200 bp are within the library size of paired-end short-read sequencing. Therefore, they may be readily detected by short-read sequencing. Second, PacBio sequencing has a fairly high per-base error rate and we found it has a very low precision on detection of small SVs from coverage data sets. Therefore, we believe that the advantage of PacBio sequencing may be the detection of large SVs that are more than 200 bp. The number of SVs in the high-confidence sets is shown in Table 2.

**Table 2.** Number of calls in the high-confidence SV sets

| Genome | Platform | Number of Deletions<br>($\geq 200\text{bp}$) | Number of Insertions<br>($\geq 200\text{bp}$) | Reference |
| --- | --- | --- | --- | --- |
| NA12878 | Illumina | 2094 | 1114 | [17, 18] |
| HX1 | PacBio | 2387 | 2937 | [20] |

### Performance Evaluation of SV callers

The SV calls of each caller were compared with the high-confidence SV set. Precision, recall, and F1 score were used to evaluate the performance of the callers. Precision, recall, and F1 were calculated as

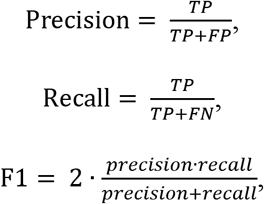
where TP is the number of true positives (variants called by a variant caller and matching the high-confidence set), FP is the number of false positives (variants called by a variant caller but not in the high-confidence set), and FN is the number of false negatives (variants in the high-confidence set but not called by a variant caller).

## Results

### Performance of SV calling on different coverages of the NA12878 genome

To determine the optimal coverage for SV detection on PacBio data, we evaluated the performance of NextSV under several different coverages. We downloaded a recently published PacBio data set of NA12878 [26] and down-sampled the data set to 2X, 4X, 6X, 8X, 10X, 12X, and 15X. SV calling was performed using NextSV under each coverage. We performed five subsampling replicates for each coverage so that the down-sampling errors could be estimated. All supported aligner/SV caller combinations were evaluated. At least two supporting reads was required for all SV calls. The resulting calls were compared with the high-confidence SV set (including 2094 deletion calls and 1114 insertion calls) described in the Method section.

First, we examined how many calls in the high-confidence set can be discovered. As shown in Figure 2, the recall increased rapidly before 10X coverage but the slope of increase slowed down after 10X. The standard deviations of recall values of the down-sampling replicates were very small (shown as error bars in the Figure). Among the four aligner / SV caller combinations, BLASR / PBHoney-Spots had the highest recall for insertions while NGMLR / Sniffles had the highest recall for deletions. At 10X coverage, BLASR / PBHoney-Spots had an average recall of 76.2% for deletions and an average recall of 81.5% for insertions; NGMLR / Sniffles had an average recall of 91.1% for deletions and an average recall of 76.3% for insertions. BWA / Sniffles had a lower recall for deletions (72.6%) and insertions (50.8%) than NGMLR / Sniffles, indicating that NGMLR was a better aligner for Sniffles. PBHoney-Tails only detected 26.3% deletions and 0.1% insertions. NextSV sensitive call set, which was generated by the union call set of BLASR / PBHoney-Spots, BLASR / PBHoney-Tails, and NGMLR / Sniffles, had the highest recall. At 10X coverage, the average recall of NextSV sensitive call set is 94.7% for deletions and 87.8% for insertions. At 15X coverage, the recall of NextSV sensitive call set increased slightly. Therefore, 10X coverage might be an optimal coverage to use in practice, considering the relatively high sequencing costs and the generally high recall rates.

**Figure 2.**
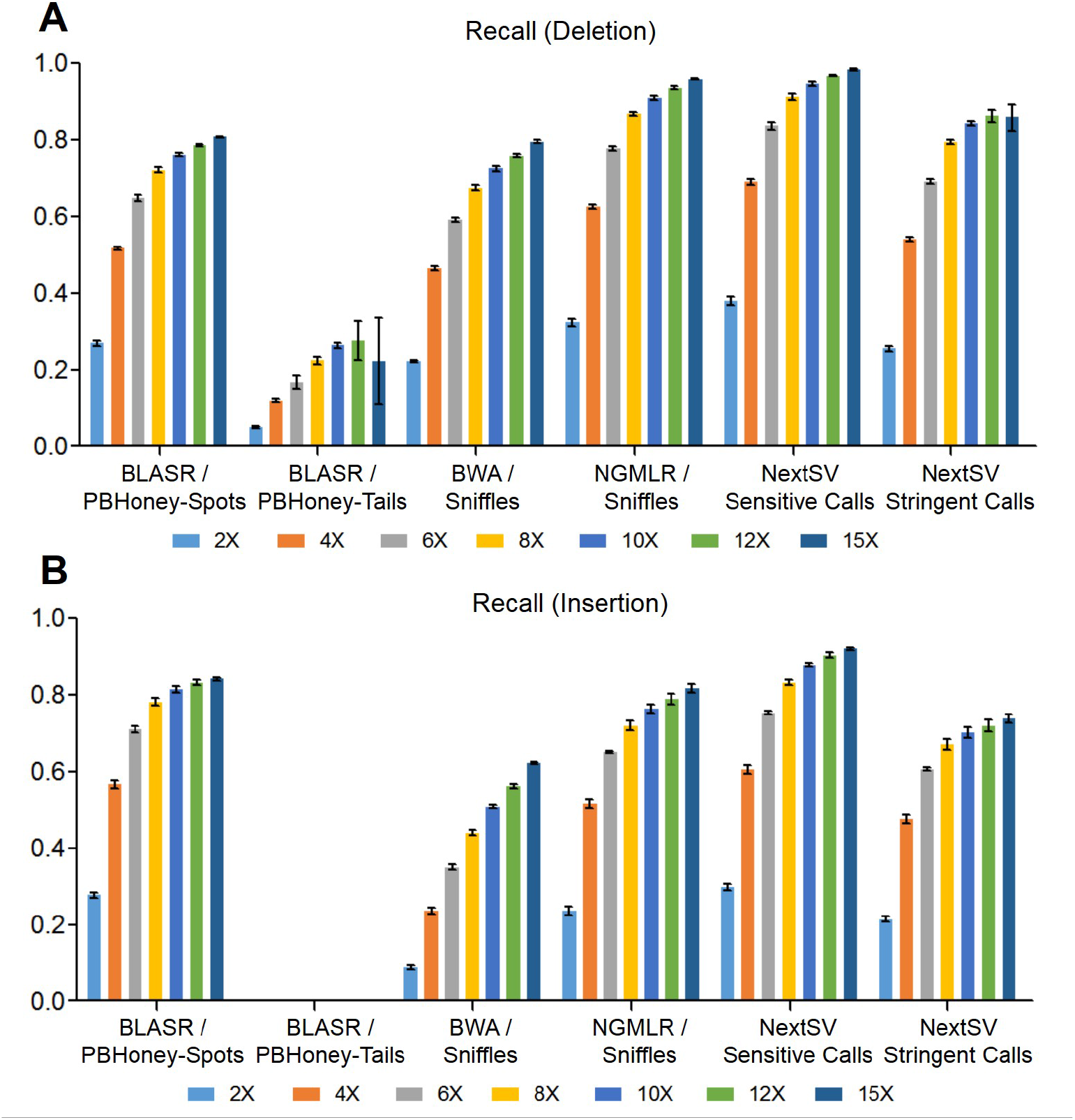
Evaluation of recall rates under different coverages on the NA12878 genome. Five down-sampling replicates were performed at each coverage. (A) Recall rates of deletion calls. (B) Recall rates of insertion calls. Data shown represent mean ± SD.

Second, we examined the precision and balanced accuracy (F1 scores) under different coverages (Figure 3). The precision was calculated as the fraction of detected SVs which matching the high-confidence set. For deletions calls, NextSV stringent call set had the second highest precision and highest F1 score. For insertion calls, NextSV stringent call set had the highest precision and F1 score at each coverage. Therefore, NextSV stringent call set performs the best, considering the balance between recall and precision. We observed that the precision decreased as the coverage increased from 2X to 15X. This was because we used the same parameter (at least two supporting reads) to generate the calls for each coverage. Therefore, the false positive rates increased as the coverage increased. A stricter parameter (e.g. at least three supporting reads) for 10X and 15X coverages may increase the precision, but decrease the recall. We discussed the trade-off between recall and precision in the Discussion section. Detailed values of recall rates, precisions and F1 scores on different coverages of the NA12878 genome were shown in Table S1-S12.

**Figure 3.**
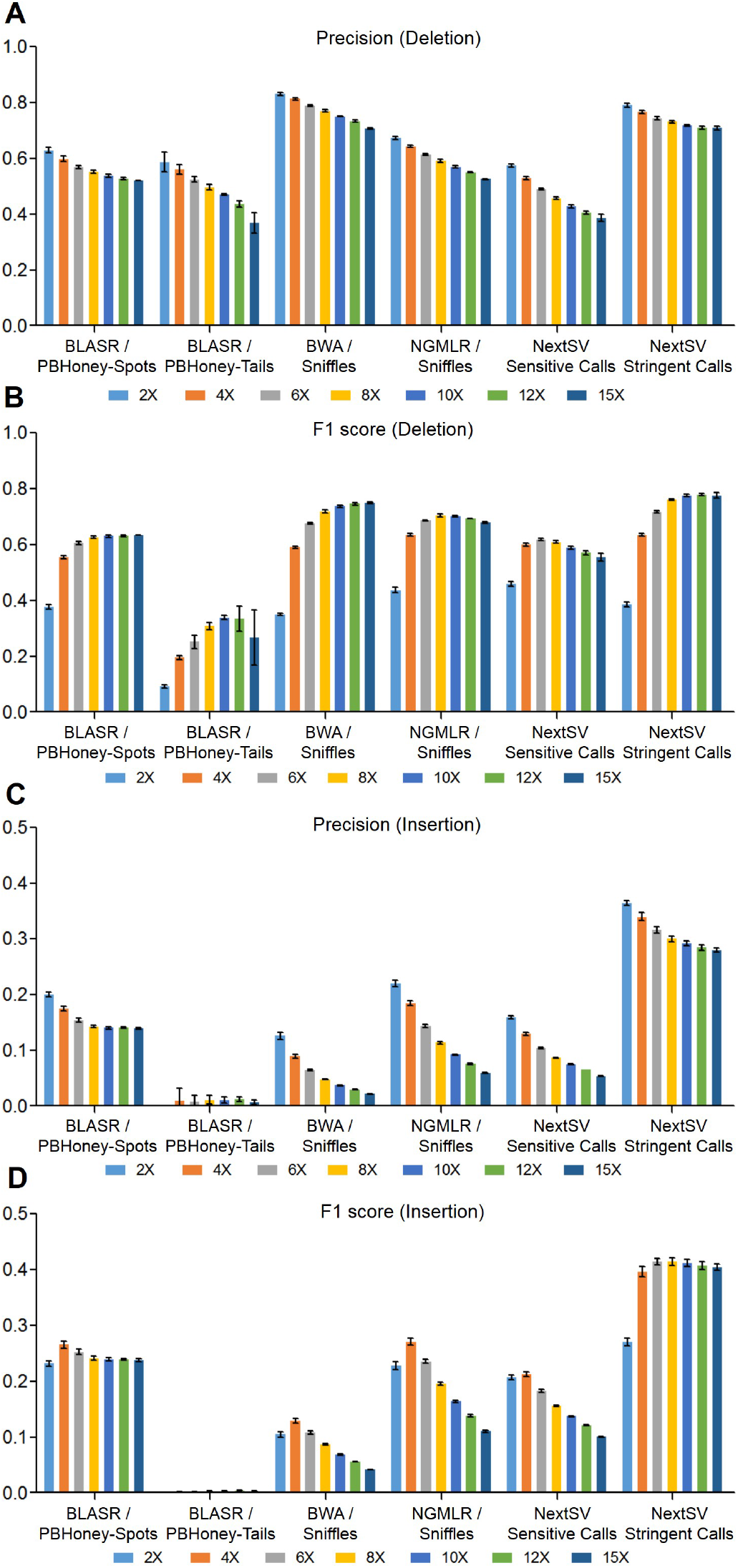
Evaluation of precisions and F1 scores under different coverages on the NA12878 genome. Five down-sampling replicates were performed. (A) Precisions of deletion calls. (B) F1 scores of deletion calls. (C) Precisions of insertion calls. (D) F1 scores of insertion calls. Data shown represent mean ± SD.

### Performance of SV calling on different coverages on the HX1 genome

To verify the performance of SV detection on different individuals, we also performed evaluation on a Chinese genome HX1, which was sequenced by us recently [20] at 103X PacBio coverage. The genome was sequenced using a newer version of chemical reagents and thus the mean read length of HX1 was 40% longer than that of NA12878 (Table 1). The total data set was downsampled to three representative coverages (6X, 10X and 15X). We also performed five subsampling replicates at each coverage. SVs were called using the four pipelines described above and compared to the high-confidence set. The results were similar to those of the NA12878 data set (Figure 4). At 10X coverage, NextSV sensitive call set had an average recall of 95.5% for deletions and 90.3% for insertions, highest among all the call sets. NextSV stringent call set had the highest precisions and F1 scores. Among the four aligner / SV caller combinations, NGMLR / Sniffles discovered the most deletions (91.6%) and BLASR / PBHoney-Spots discovered the most insertions (81.5%) at 10X coverage. BWA / Sniffles had a higher precision but a lower recall and F1 score than NGMLR / Sniffles. Detailed values of recall rates, precisions and F1 scores on different coverages of the HX1 genome were shown in Table S13-S24.

**Figure 4.**
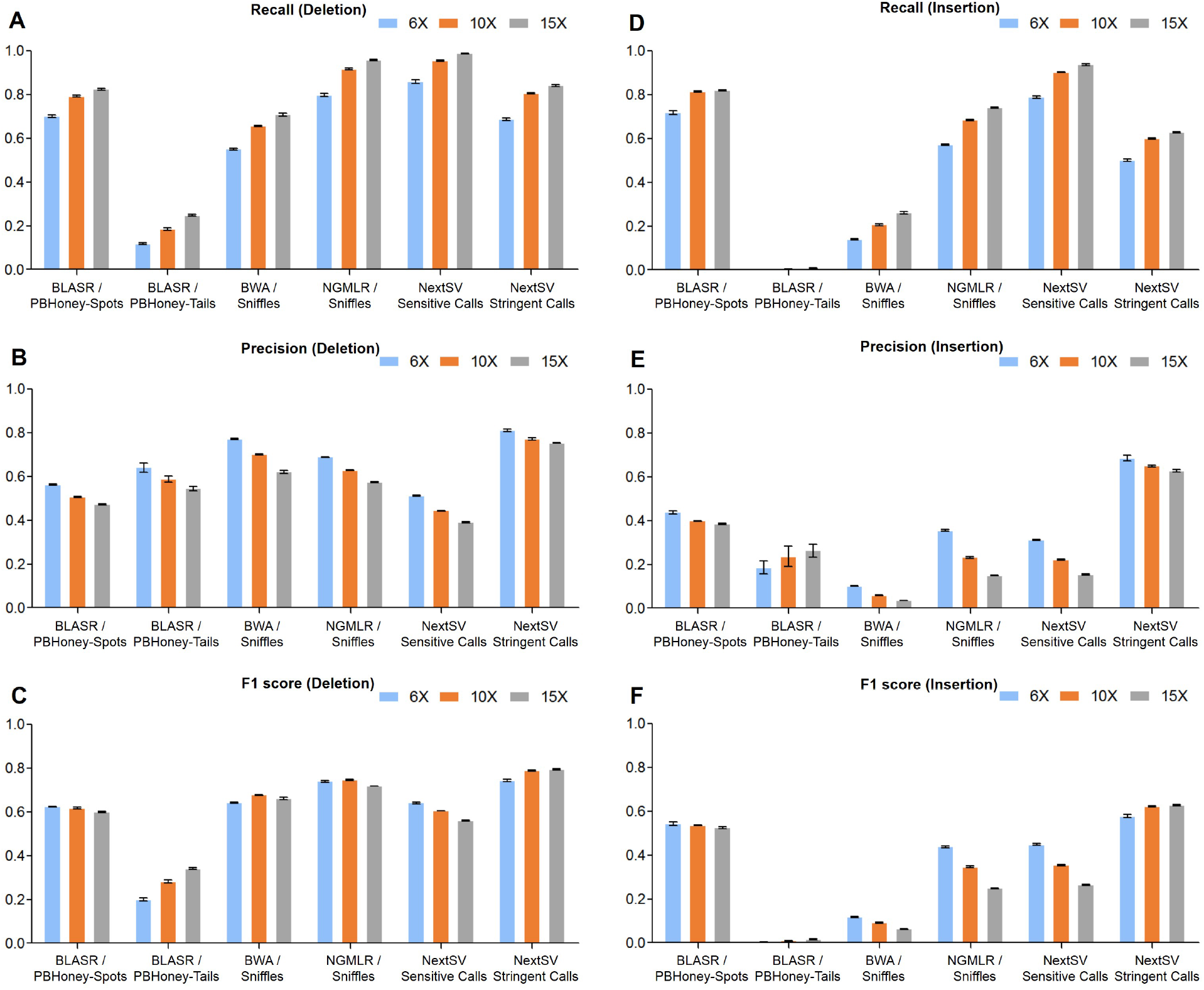
SV calling performance on the HX1 genome. Five down-sampling replicates were performed. (A-C) Recall rates, precisions and F1 scores of deletion calls. (D-F) Recall rates, precisions and F1 scores of insertion calls. Data shown represent mean ± SD.

### Evaluation on Mendelian Errors

As the *de novo* mutation rate is very low [27, 28], Mendelian errors are more likely a result of genotyping errors and can be used as a quality control criteria in genome sequencing [29]. Due to the lack of gold standard call sets, here we evaluated the errors of allele drop-in (ADI), which means the presence of an allele in offspring that does not appear in either parent. The ADI rate is calculated as the ratio of ADI events to SV calls detected in the offspring. We used a whole genome PacBio sequencing data set of an AJ family trio released by NIST [19] to do the evaluation. The sequencing data for father, mother and son are 32X, 29X, and 63X, respectively. First, we did the ADI rate analysis using all the available data. Since the coverages were high, 8 supporting reads were required for SV calls of the parents and 15 supporting reads were required for SV calls of the son. Among the four aligner/SV caller combinations, NGMLR/Sniffles had the lowest ADI rate (12.0%) for deletions, while BLASR/PBHoney-Tails had the lowest ADI rate (10%) for insertions (Figure 5). Next, we down-sampled the sequencing data of the son to 10X coverage and analyzed the ADI rate at this low coverage. Five down-sampling replicates were performed. The ADI rates at 10X coverage were generally higher than those at 63X coverage. NGMLR/Sniffles achieved lowest ADI rate for both deletions (19.0%) and insertions (25.2%) among the four aligner/SV caller combinations. NextSV stringent call set had the lowest ADI rate for insertions (15.7%) and second lowest ADI rate for deletions (20.0%). The standard deviations of ADI rates of the downsampling replicates were very small (shown as error bars in the Figure).

**Figure 5.**
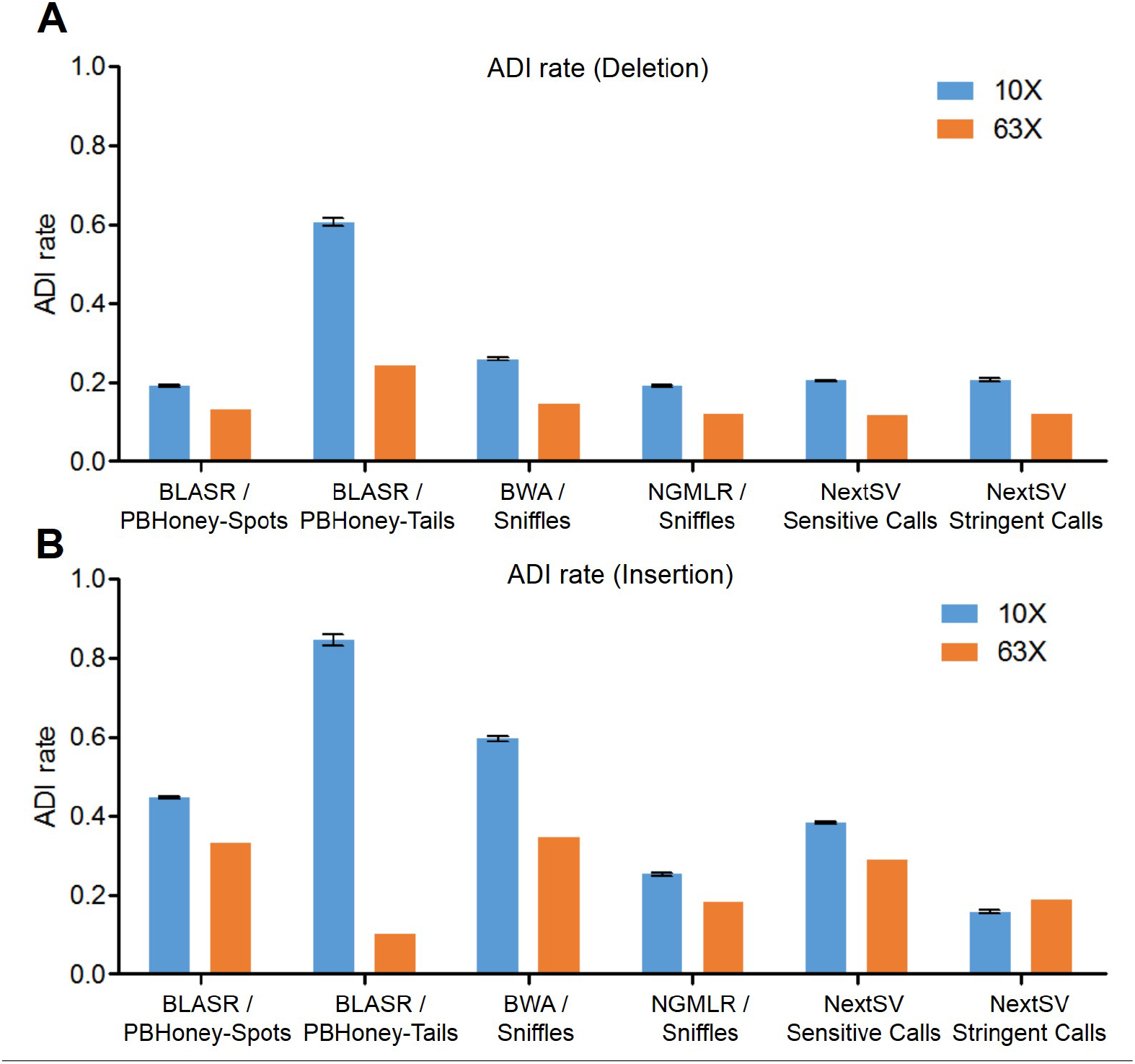
Comparison of allele drop-in rate. For evaluation of ADI rate at 10X coverage, five down-sampling replicates were performed. (A) ADI rates of deletion call. (B) ADI rate of insertion calls. Data shown represent mean ± SD.

### Computational Performance of NextSV

To evaluate the computational resources consumed by NextSV, we used the whole genome sequencing data set of HX1 (10X coverage) for benchmarking. All aligners and SV callers in NextSV were tested using a machine equipped with 12-core Intel Xeon 2.66 GHz CPU and 48 Gigabytes of memory. As shown in Table 3, mapping is the most time-consuming step. BLASR takes about 80 hours to map the reads, whereas NGMLR needs only 11.2 hours. The SV calling step is much faster. PBHoney-Spots and Sniffles take about 1 hour, while PBHoney-Tails needs 0.27 hour. In total, the BLASR / PBHoney combination takes 80.8 hours while the NGMLR / Sniffles combination takes 12.5 hours, 84.5% less than the former one. Since BLASR/PBHoney-Spots and NGMLR / Sniffles have good performance on SV calling and running PBHoney-Tails is very fast given the BLASR output, the NextSV pipeline will execute the three methods by default for generating the final results.

**Table 3.** Time consumption for each steps in the NextSV pipeline for 10X PacBio data set

| SV caller | Aligner | CPU<br>(number of threads) | Alignment time<br>(hour) | SV calling time<br>(hour) | Total Time<br>(hour) |
| --- | --- | --- | --- | --- | --- |
| PBHoney | BLASR | 12 | 79.6 | 0.27 (Tails)<br>0.96 (Spots) | 80.8 |
| Sniffles | BWA-MEM | 12 | 27.0 | 1.1 | 28.1 |
| Sniffles | NGMLR | 12 | 11.2 | 1.3 | 12.5 |

## Discussion

Long-read sequencing such as PacBio sequencing has clear advantages over short-read sequencing on SV discovery [10]. However, its application in real-world setting is often limited due to the relatively high sequencing cost and hence the relatively low sequencing coverage. Some efforts have been made to improve SV detection from low coverage short-read data[30], but methods for improving SV detection from long-read sequencing data have not been reported. In this study, we developed NextSV, a meta SV caller integrating multiple aligners and SV callers to improve SV discovery on low-coverage PacBio data sets. Our results showed that, NextSV stringent call set had the highest precisions and F1 scores while NextSV sensitive call set had the highest recall. At 10X coverage, the recall of NextSV sensitive call set was 94.7% to 95.5% for deletions and 87.8% to 90.3% for insertions. At 15X coverage, there was only a slight increase in recall. Therefore, ~10X coverage can be an optimal coverage to use in practice, considering the balance between the sequencing costs and the recall rates.

The high-confidence call set of HX1 genome was generated using two steps. First, we used a call set from a previously validated local assembly-based approach [11, 20, 31] as the initial high-quality calls. Second, we detected SVs on 103X coverage PacBio data set of the HX1 genome using the four aligner/SV caller combinations described above. The calls were filtered using a strict parameter (minimal read support=20 for each SV caller). The initial high-quality calls that overlapped with one of the four 103X call sets were retained as final high-confidence calls. Since the aligners/SV callers contribute to generation of the high-confidence call sets, there may be some biases on the comparison of aligner/SV callers. However, it would be less biased on comparison of the performances on different coverages, which is an important goal of our study.

There is often a trade-off between recall and precision. NextSV generates a sensitive call set and a stringent call set, for different purposes. NextSV sensitive call set is suitable for users who consider recall more important than precision and who can afford extensive downstream analysis (such as Sanger sequencing) to validate the candidate variants. This is often the case when doing disease-casual variant discovery on personal genomes. NextSV stringent call set has the highest precision, F1 score. It is suitable for users who aim to perform genome-wide analysis of SVs on a collection of samples, with limited downstream validation.

The performance of SV callers are affected by the parameter settings. The number of supporting reads is a key parameter that affect the trade-off between recall and precision. By default, PBHoney requires a minimal read support of 3 for an SV event and Sniffles requires a minimal read support of 10 for an SV event. However, this may be too high for low coverage data set. In our evaluation of recall and precision, we changed this setting to require a minimal read support of 2. This allows detection of SVs from very low coverage regions, with an acceptable precision. Thus, substantially higher number of true positives would be detected and less variants of interest would be missed. For users who consider precision to be more important than recall, they can either use the NextSV stringent call set or specify a stricter parameter (e.g. requiring more supporting reads) when running the NextSV pipeline. The F1 score is a balance between recall and precision. Therefore, its correlation with coverage is affected by the two aspects. In general, as the coverage increases, the recall increases but the precision decreases. Therefore, the F1 score may either increase or decrease as the coverage increases.

In addition to test recalls and precisions, we examined the allele drop-in (ADI) errors, which represent the SV calls that in the offspring but not appear in either parent. Since the *de novo* mutation rate is very low, the ADI errors may mainly come from errors of sequencing and subsequent SV detection. In our results, the ADI rates of insertions are higher than those of deletion calls, which is consistent with the fact that PacBio sequencing has higher per-base insertion errors than deletion errors. Another source of ADI may come from the SV callers. SV detection from PacBio data set is still in its early stage. The currently available SV callers are not carefully designed for low-coverage data sets. For example, Sniffles requires 10 reads to support a SV under default settings, which means at least 20X coverage is required to detect a heterozygous SV. We expect the improvement of SV callers in the future.

NextSV currently supports four aligner / SV caller combinations: BLASR / PBHoney-Spots, BLASR / PBHoney-Tails, BWA / Sniffles, NGMLR / Sniffles, but we expect to continuously expand the support for other aligner / caller combinations. In the future, if more aligners/SV callers are supported, we will evaluate the performance of each combination and find the best aligner for each SV caller. The NextSV sensitive call will be the union call set of all SV callers; the NextSV stringent calls will be the calls that are detected by at least two SV callers. If one SV caller can work with multiple aligners, only the call set of its best aligner will be used.

In this study, we only evaluated the performance for insertions and deletions because we only have the high-confidence calls of insertions and deletions. This is another limitation of the study. We will evaluate the performance on other types of SVs in the future when more high-confidence SV calls are available. Nonetheless, NextSV generates SV calls of all types. The output of NextSV is in ANNOVAR-compatible format. Users can easily perform downstream annotation using ANNOVAR and disease gene discovery using Phenolyzer [32]. NextSV is available on GitHub [33] and can be installed by one simple command.

## Conclusion

In this study, we proposed NextSV, a comprehensive, user-friendly and efficient meta-caller to perform SV calling from low coverage long-read sequencing data. NextSV integrates multiple aligners and SV callers and performs better than running a single SV caller. We also showed that ~10X PacBio coverage can be an optimal coverage to use in practice, considering the balance between the sequencing costs and the recall rates. Our results provide useful guidelines for SV detection from low coverage whole-genome PacBio data and we expect that NextSV will facilitate the analysis of SVs on long-read sequencing data.

## Abbreviations

SV: structural variant; NGS: next-generation sequencing; SMRT: single-molecule, real-time; GIAB: Genome in a Bottle; NIST: National Institute of Standards and Technology; AJ: Ashkenazi Jewish; ADI: allele drop-in

## Author’s Contributions

LF performed the evaluation, implemented the software and wrote the manuscript. JH and DW tested the software and advised on the study. DW and KW conceived and supervised the study, and revised the manuscript. All authors read and approved the final manuscript.

## Acknowledgments

The authors wish to thank the National Institute of Standards and Technology and Genome in a Bottle Consortium for making the reference data on PacBio sequencing available to benchmark bioinformatics software tools. We also thank members of Grandomics to test the software tools and offering valuable feedback.

## Availability of Data and Materials

The PacBio sequencing data of NA12878 and HX1 analyzed in this study are available in the NCBI SRA database (Accession: SRX627421, SRX1424851). The PacBio sequencing data of AJ trio family is available at the FTP site of NIST (ftp://ftp-trace.ncbi.nlm.nih.gov/giab/ftp/data/AshkenazimTrio/, release date: Nov 9^th^, 2015). NextSV is available at http://github.com/Nextomics/NextSV.

## Ethics approval and consent to participate

Not applicable.

## Consent for publication

Not applicable.

## Supplemental file

Supplemental file 1: Tables S1-S24. Performances of BLASR/PBHoney-Spots, BLASR/PBHoney-Tails, BWA/Sniffles, NGMLR/Sniffles and NextSV on the NA12878 genome and the HX1 genome. (PDF 473 kb)

